# Limited Defense of Liver Energy Status in Rats Susceptible to Diet-induced Obesity

**DOI:** 10.1101/2022.10.26.513713

**Authors:** Mark I Friedman, Hong Ji

**Affiliations:** Monell Chemical Senses Center, Philadelphia, Pennsylvania, United States of America

**Keywords:** obesity, liver, ATP, fasting, food intake

## Abstract

Rats that are susceptible to diet-induced obesity have a preexisting reduced capacity for hepatic fatty acid oxidation compared with those resistant to diet-induced obesity. The eating response to administration of a fatty acid oxidation inhibitor is more closely associated with low liver energy status than it is with reduced hepatic fatty acid oxidation, a finding consistent with studies showing that lowered liver energy status stimulates food intake. To evaluate whether susceptibility to diet-induced obesity is associated with a preexisting impairment in liver energy status, we conducted two experiments in obesity-prone (OP) and -resistant (OR) outbred rats. In one experiment, OP rats increased food intake more than did OR rats during refeeding after a 24 h fast. When fasted and refed again, liver energy status (i.e., liver ATP content, ATP:ADP ratio and phosphorylation index) was lower in OP rats after a fast. When OP animals were refed fixed rations of food, liver energy status increased more slowly during refeeding than it did in OR rats. The delay in restoration of liver energy status during refeeding corresponded to the interval during which OP rats ate significantly more food in the intake test. In a second experiment, liver energy status was lower in OP than it was in OR rats after injection of the fructose analogue, 2,5-anhydro-D-mannitol, which depletes liver ATP. These results suggest that liver energy status is more vulnerable in rats susceptible to diet-induced obesity.

## Introduction

Diet composition is considered an important contributing factor to the development of obesity. In animal models of diet-induced obesity, diets containing substantial amounts of both carbohydrate and fat produce the greatest weight gain and increase in energy intake (1,2). There is a high degree of individual variability in weight gain among laboratory rodents fed such diets; while some animals become obese, other remain lean (3–5). Susceptibility to diet-induced obesity in rats appears to be tied to a reduced capacity for hepatic fatty acid oxidation that is evident prior to feeding a high-carbohydrate/high-fat (HC/HF) diet (6–9). However, the eating response to administration of a fatty acid oxidation inhibitor in lean rats is more closely associated with reduced liver energy status (i.e., liver ATP content, ATP:ADP ratio and phosphorylation index) than it is with reduced liver fatty acid oxidation (10). This finding raises the possibility that differences in hepatic energy production, more so than that in fat oxidation, may underlie susceptibility to diet-induced obesity.

Numerous studies suggest that changes in liver ATP content generate a signal to control food intake (for a review see 11). Much of the direct evidence stems from experiments in rats using the fructose analogue, 2,5-anhydro-D-mannitol (2,5-AM), which reduces liver ATP by trapping phosphate in its phosphorylated forms, which are not further metabolized (12). Administration of 2,5-AM stimulates food intake by acting in the liver (13) to trap phosphate and reduce ATP (14,15). 2,5-AM administration also increases whole body and hepatocyte fatty acid oxidation (16,17), which appears to restrain the eating response. Increasing fat oxidation by feeding a high-fat/low-carbohydrate diet attenuates the reduction in hepatocyte and liver ATP and prevents the eating response to 2,5-AM (17,18). Co-administration of 2,5-AM and a methyl palmoxirate, a fatty acid oxidation inhibitor, when each is given in a dose that alone has no effect, synergistically decreases liver energy status and increases food intake (19). This latter finding suggests that reduced fatty acid oxidation can limit the capacity of the liver to maintain its energy status.

We performed two experiments in an initial effort to investigate whether an impaired ability to defend liver energy status predisposes rats to diet-induced obesity. Fasting reduces liver ATP in rats (20,21) and liver energy status (as measured by ATP, ATP:ADP ratio and/or phosphorylation index) is restored after animals are refed with a time course that parallels the compensatory hyperphagia during refeeding (21). In one experiment, liver energy status was assessed in obesity-prone (OP) and -resistant (OR) rats after fasting and during refeeding. A second experiment compared the effects of 2,5-AM injection on liver energy status in OP and OR rats.

## METHODS AND PROCEDURES

### Animals and diets

Male Sprague-Dawley CD rats from Charles River Laboratory (Wilmington, MA) were housed individually in hanging stainless steel cages in a vivarium maintained at 22°C and a 12:12 h day:night cycle. Rats had access to bottled tap water and food *ad libitum* unless otherwise noted. All animal protocols were in compliance of NIH guidelines for animal care and use, and approved by the Institutional Animal Care and Use Committee of the Monell Chemical Senses Center.

Three diets were used in these experiments: (i) a standard rodent chow (LabDiet #5001); (ii) a semi-synthetic high-carbohydrate/low-fat (HC/LF) diet containing, by calories, 13% fat, 63% carbohydrate and 24% protein with an energy density of 3.3 kcal/g; (iii) a semi-synthetic high-energy, high-carbohydrate/high-fat (HC/HF) diet containing, by calories, 42% fat, 41% carbohydrate, 17% protein with an energy density of 4.7 kcal/g. The semi-synthetic diets were custom-made by Dyets, Inc. (for a compositions see 1).

### Tissue and Plasma Analyses

Rats were anesthetized by an intramuscular injection of Ketamine HCl (100 mg/kg) plus acepromazine maleate (1 mg/kg). Via a midline abdominal incision, a sample of the median lobe of the liver was excised and freeze-clamped immediately using a pair of aluminum blocks pre-chilled in liquid nitrogen. Trunk blood was collected into a chilled centrifuge tube containing EDTA and Trasylol®. Plasma was prepared within 30 minutes from the blood chilled on ice. Liver and plasma samples were stored at -80°C until being analyzed, usually within 2 wks.

Frozen liver samples were extracted with HClO4 and analyzed for ATP and ADP by HPLC (14). Inorganic phosphate (Pi) in the liver extracts was determined using the Sigma Kit 360-3. Phosphorylation potential was calculated as [ATP]/([ADP] × [Pi].

Plasma concentrations of glucose and triglyceride were determined using Sigma kits. Total ketone bodies (β-hydroxybutyrate and acetoacetate) and glycerol were analyzed fluorometrically (22). Free fatty acids (FFA) were determined using a NEFA Kit from Wako Chemicals USA, Inc. Insulin, leptin and glucagon were quantified by the RIA/Biomarkers Core at the University of Pennsylvania.

### Data analysis

Time course data for OP and OR groups were compared using two-way ANOVA with repeated measures as appropriate. Individual post-hoc comparisons were made using a Tukey test or t-test as appropriate.

### Experiment 1. Food intake and liver energy status during refeeding after a fast

This experiment was designed to examine whether OP and OR rats differ with respect to the degree of compensatory hyperphagia displayed after fasting and whether any such differences are reflected in changes in liver energy status or other metabolic parameters before and during refeeding.

Seventy-two rats (150-175 g) were fed chow upon arrival into the laboratory. On the next day rats were weighed and fed the HC/HF diet for 1 wk and reweighed. Twenty-four rats each with the greatest and least weight gains were designated as OP and OR respectively. Previous experiments indicated that a 1-wk weight gain of rats fed the HC/HF diet was predictive of longer term (2-4 wk) weight gain (6). OP and OR rats were then given the HC/LF diet for 2 wks to reverse the diet-induced weight gain. Rats were then deprived of food at the onset of dark period and 24 h later were refed the HC/HF diet and intakes measured 2, 6 and 12 h afterwards.

Rats were then fed the HC/LF diet *ad libitum* again for 1wk at which time the groups of OP and OR rats were each divided into 4 groups of 6 animals matched for body weight within each group, and fasted for 24 h as before. One group each of OP and OR rats were then anesthetized for tissue and blood collection (0 h). Food intake in the first 2 h of the *ad libitum* refeeding test did not differ significantly between the two groups (5.8 and 5.5 g for, respectively, OP and OR rats). Therefore, to assess metabolic responses to refeeding, rats were refed a fixed amount of the HC/HF diet equal to 75% of the average intake by OP and OR rats during the first 2 h of refeeding during the intake test the previous week to control for differences in intake between groups and assure that rats consumed all their food within the first 2 h of the test to avoid effects of ongoing food consumption on metabolic measures. One group each of OP and OR rats was then anesthetized for tissue and blood collection 3, 6 and 12 h during the refeeding period.

### Experiment 2. Effect of 2,5-AM on hepatic energy status in OP and OR rats

To avoid any confounding effects of differences in weight gain after feeding the HC/HF diet to identify OP and OR rats (see results), we screened chow-fed rats for susceptibility to diet-induced obesity as described previously (9) by measuring changes in plasma triglycerides from baseline after an intragastric injection of a corn oil and Polycose mixture. Twenty-four rats (125-150 g) fed chow for 1 wk after arrival into the laboratory were thus screened and 8 rats with the smallest changes in plasma TG levels were identified as OP and another eight with the greatest changes were identified as OR. These 16 rats were given the HC/LF diet *ad libitum for 9* days and then injected (i.p.) with 300 mg/kg 2,5-AM (Toronto Research Chemicals). Food was removed after the injection and liver samples collected 1 h later.

## RESULTS

### Experiment 1. Food intake and liver energy status during refeeding after a fast

#### Body weight

Body weights (mean + SEM) prior to feeding the HC/HF diet for OP and OR rats were, respectively, 176 ± 1 and 171 ± 1 g [*t*(46) = 2.6, *P* = 0.01]. After consuming the HC/HF diet for 1 wk, rats designated as OP, compared with OR-designated rats, gained significantly more weight [75 ± 1 vs. 56 ± 1 g; *t*(46) = 12.7, P < 0.0001] and weighed significantly more [251 ± 2 vs. 227 ± 2 g: *t*(46) = 9.4, *P* < 0.0001]. Two wks after their diet was switched from the HC/HF to the HC/LF diet, just prior to the fasting-refeeding test, OP and OR rats weighed, respectively, 356 ± 4 and 312 ± 3 g [*t*(46) = 9.7, *P* < 0.0001]. The following week when rats were retested to assess the metabolic response to fasting and refeeding, OP and OR rats weighted, respectively, 401 ± 5 and 346 ± 3 g [*t*(46) = 9.0, P < 0.0001].

#### Food intake

After 24 h of fasting, OP rats ate more of the HC/HF diet than did OR rats during the refeeding period [*F*(1,46) = 14.6, *P* = 0.0004; Figure 1]; total cumulative food intakes during the 12-h period were greater in OP than OR rats [24.7 ± 0.5 vs. 21.9 ± 0.5 g; t(46) = 3.8, P = 0.0004]. These differences in food intake were largely attributable to greater food intake of OP rats between 2 and 6 h of refeeding [5.8 ± 0.3 g and 7.1 ± 0.4 g; t(46) = 2.7, P = 0.01].

**Figure 1.**
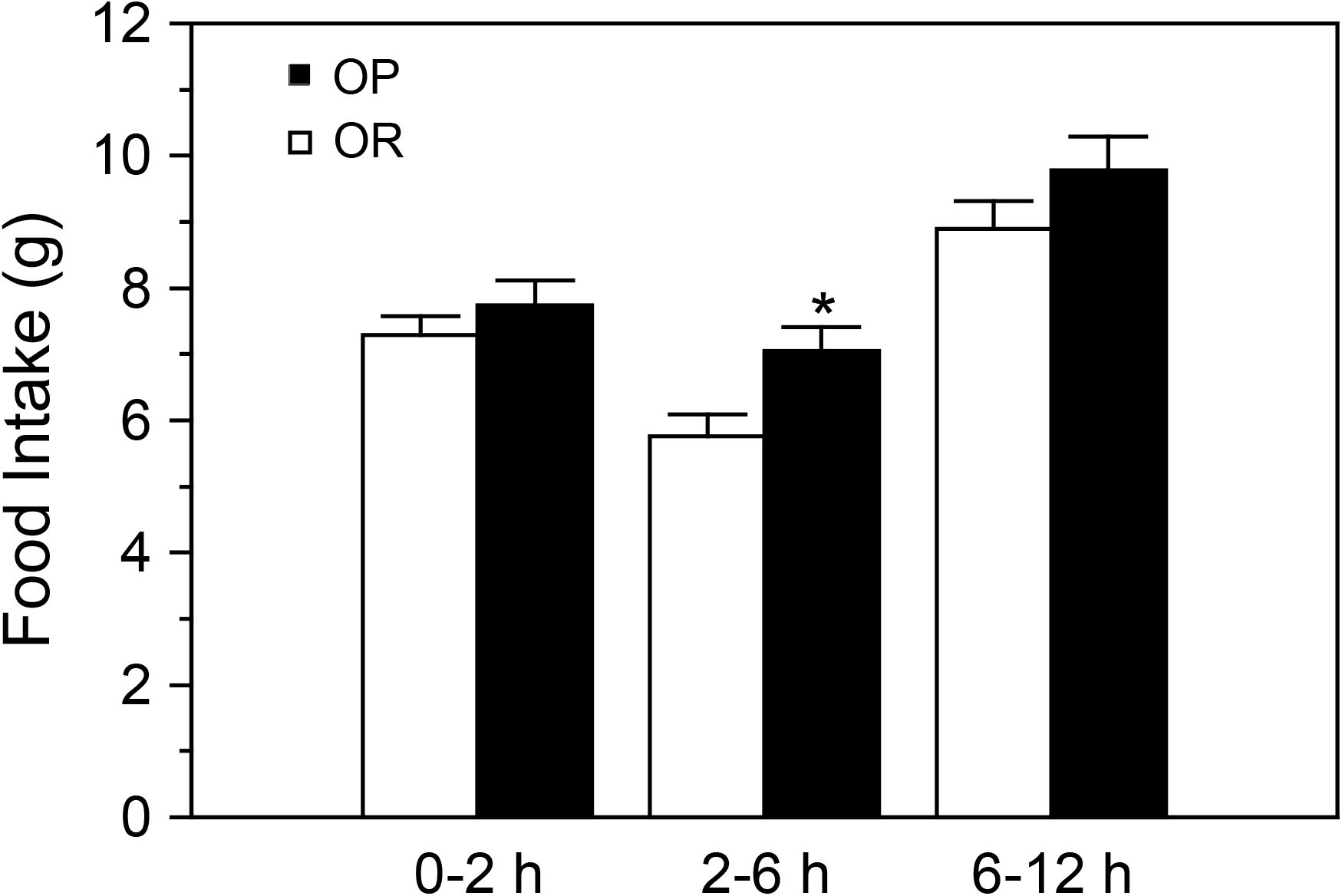
Food intake after 24 h fasting in obesity-prone (OP) and obesity-resistant (OR) rats (Ns = 12). An obesity promoting diet with approximately equal amount of calories from carbohydrate (42%) and fat (41%) was given at 0 h. Values are mean ± SEM. * denotes a significant difference (see text) between the groups.

#### Liver energy status

As shown in Figure 2, liver energy status, as measured by ATP concentration, ATP:ADP ratio, and phosphorylation potential, was lower in OP rats than it was in OR rats [*F*s(1,40) = 6.5, 16.8 and 23.6, *P*s < 0.02, 0.0003 and 0.0001 for ATP, ATP:ADP ratio and phosphorylation potential, respectively]. After fasting, liver ATP contents were similar in OP and OR rats, but ATP:ADP ratio and phosphorylation potential were lower in OP rats (*P*s < 0.01). By 3 and 6 h of refeeding, all three measures of energy status were lower in OP rats (*P*s < 0.01).

**Figure 2.**
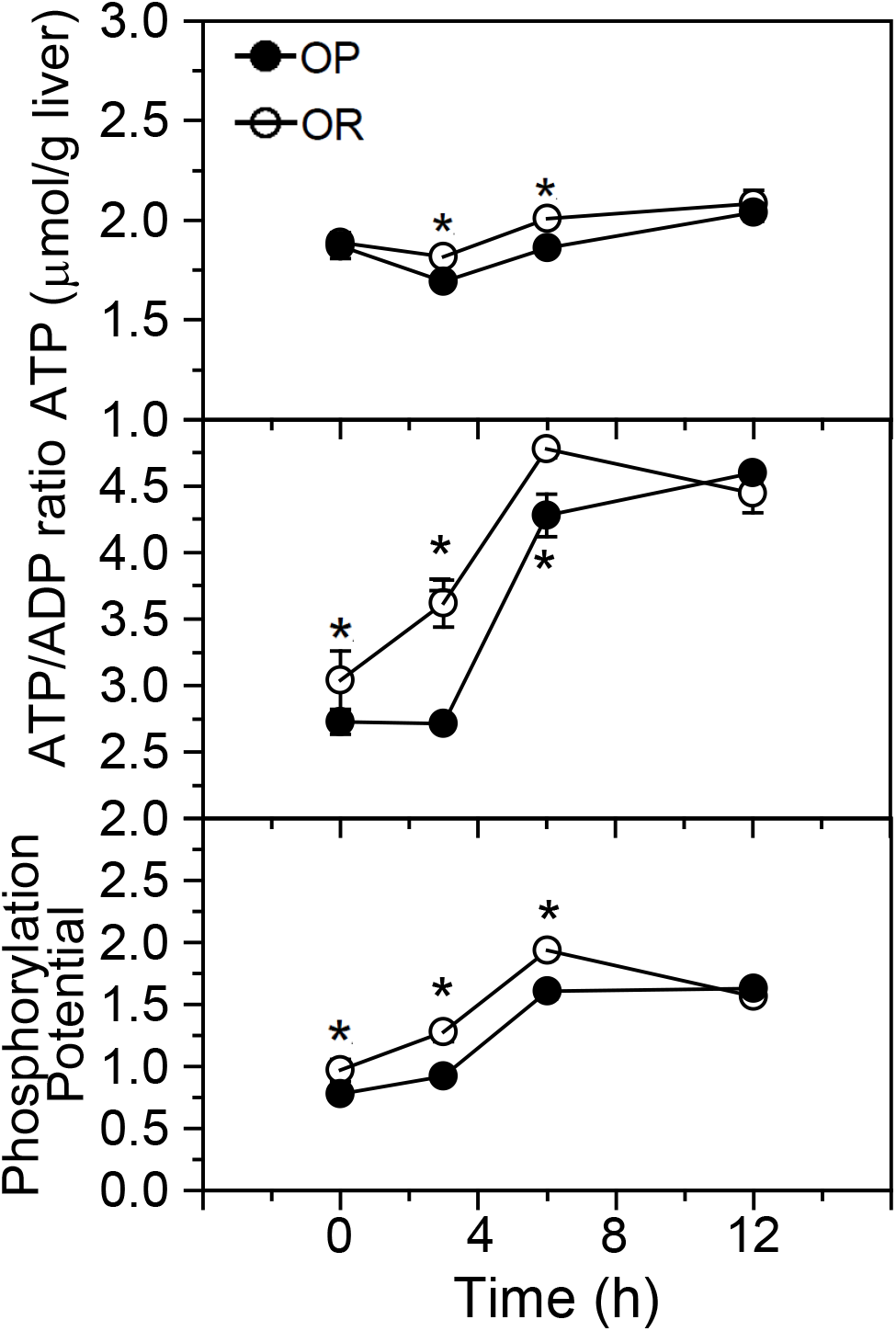
Liver energy status during restricted refeeding after 24 h fasting in obesity-prone (OP) and obesity-resistant (OR) rats (N = 6 for each group at each time point). An obesity promoting diet with approximately equal amount of calories from carbohydrate (42%) and fat (41%), and restricted to 75% of the amount eaten under *ad libitum* refeeding conditions, was given at 0 h. Values are mean ± SEM. * denotes a significant difference (see text) between the groups at the time points indicated.

Liver energy status increased in both OP and OR rats during refeeding [*F*s(3,40) = 15.8, 83.6 and 101.0, respectively for the ATP concentration, ATP:ADP ratio, and phosphorylation potential of the three measures, *P*s < 0.0001]; however, liver energy status increased more slowly in OP rats compared with OR rats. Whereas ATP concentrations in OR rats increased from fasting levels (0 h) during the first 6 h (p < 0.05), they decreased from fasting levels in OP rats during the first 3 h (P < 0.01) and did not increase above fasting levels until 12 h (*P* < 0.01) of refeeding. ATP:ADP ratio and phosphorylation potential increased more slowly over time after refeeding in OP rats than they did in OR rats [*F*s(3,40) = 5.2 and 5.3, *P*s = 0.0041 and 0.0036 for interactions for ATP:ADP ratio and phosphorylation potential, respectively]. In OR rats, the liver ATP:ADP ratio and phosphorylation potential increased from fasting levels starting by 3 h of refeeding and continuing to rise during the next 3 h (*P*s <.01). In contrast, neither measure changed from fasting levels in OP rats until 6 h of refeeding. All measures of liver energy status were similar in OP and OR rats by 12 h of refeeding.

#### Metabolic hormones

Plasma insulin concentrations (Figure 3, left) did not significantly differ over time at the sampling intervals used; nor did they differ between groups, although there was a tendency for insulin concentrations to be greater in OP rats [*F*(1,40) = 3.0, *P* = 0.089]. Considered separately, OP rats had greater insulin concentrations after fasting (0 h) than did OR rats [*t*(10) = 2.89, *P* = 0.016]. Circulating glucagon concentrations decreased during refeeding [*F*(3,40) = 3.7, *P* = 0.019] and were overall greater in OP rats compared with OR rats [*F*(1,40) = 6.8, *P* = 0.013; Figure 3, left]. Glucagon concentrations were greater in OP as compared with OR rats after fasting (0 h; *P* < 0.05) and after 3 and 12 h of refeeding (*P*s < 0.01). Insulin:glucagon ratios (data not shown) did not differ between groups, but did increase from fasting levels during refeeding [F(3,40) = 4.0, P = 0.015]. Plasma leptin concentrations were below the level of detection in one OP rat at 12 h, three OR rats taken at 0 h and four OR rats at 12 h. Plasma leptin concentrations in the remaining rats (Figure 3, left) increased from fasting levels to 6 h of refeeding and then declined back to baseline. This change over time was more pronounced in OP compared with OR rats [*F*(3,32) = 3.3, *P* = 0.034 for interaction] with OP rats showing greater leptin concentrations at all but 0 h (Ps < 0.010).

**Figure 3.**
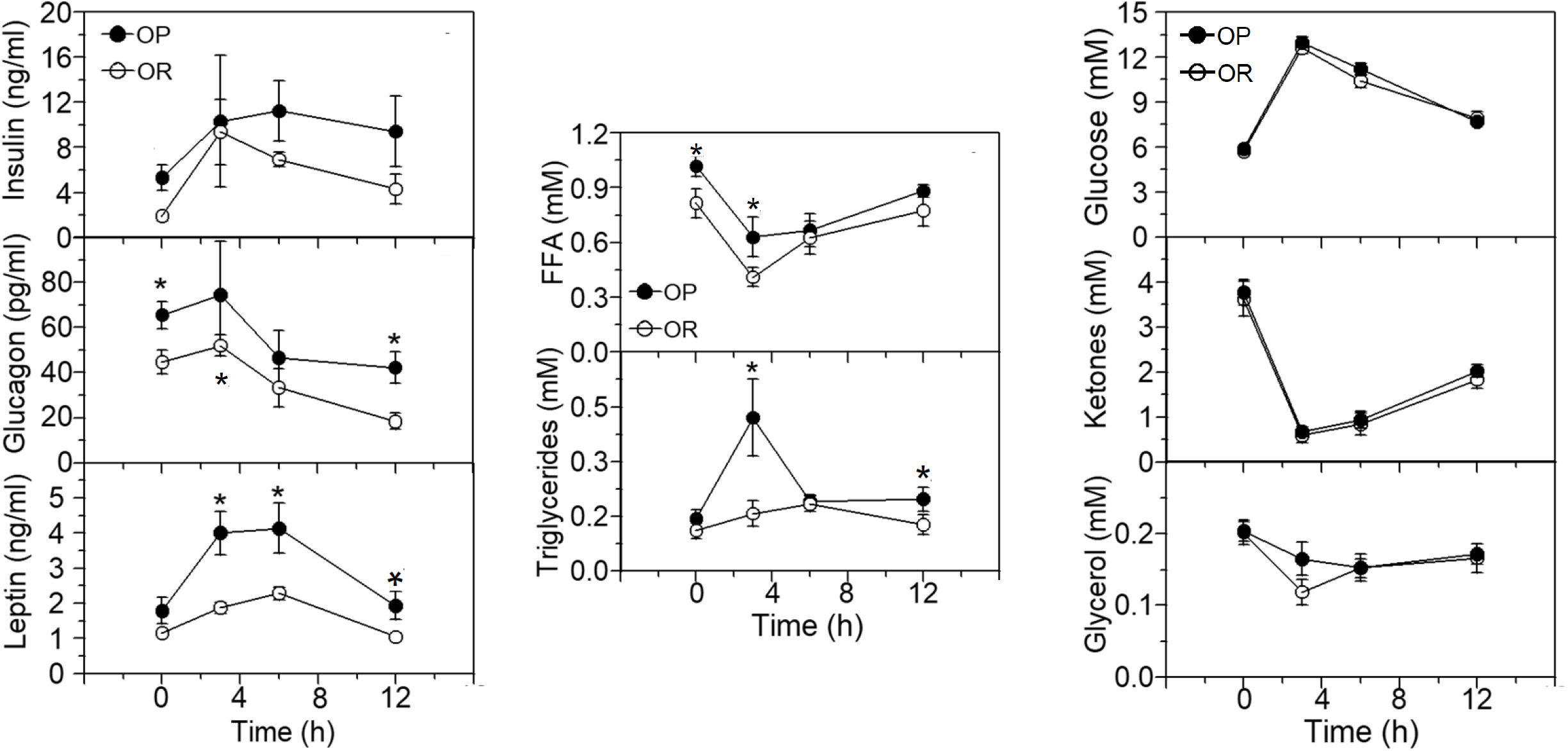
Metabolic hormones, substrates and fuels during restricted refeeding after 24 h fasting in obesity-prone (OP) and obesity-resistant (OR) rats (N = 6 for each group at each time point). An obesity promoting diet with approximately equal amount of calories from carbohydrate (42%) and fat (41%), and restricted to 75% of the amount eaten under *ad libitum* refeeding conditions, was given at 0 h. Values are mean ± SEM. * denotes a significant difference (see text) between the groups at the time points indicated.

#### Metabolic fuels and substrates

The concentration of plasma FFA (Figure 3, middle) decreased from fasting levels during the first 3 h after refeeding in both OP and OR groups of rats, and then increased to near baseline concentrations by 12 h [*F*(3,40) = 10.4, *P* < 0.0001]. Overall, plasma FFA concentrations were greater in OP rats compared to OR rats [*F*(1,40) = 6.5, *P* = 0.015] with significant differences after fasting (0 h) and after 3 h of refeeding (*P*s < 0.01). Plasma triglyceride levels increased during refeeding only in OP rats [*F*(3,40) = 3.3, *P* = 0.029 for interaction; Figure 3, middle] with significant increases 3 and 6 h after the start of refeeding [*P*s < 0.01 and 0.05, respectively]. Plasma glucose levels (Figure 3, right) were sharply greater 3 h after the start of refeeding in both groups and then decreased over the next 9 h, although not to fasting (0 h) levels [*F*(3,40) = 154.9, *P* < 0.0001]. Plasma ketone body concentrations (Figure 3, right) decreased markedly in both groups by 3 h after rats started to refeed and then rose somewhat over the next 9 h [*F*(3,40) = 74.4, *P* < 0.0001]. Glycerol concentrations in plasma (Figure 3, right) decreased slightly in both groups after the start of refeeding, and remained so for the next 9 h measurement period [*F*(3,40) = 5.7, *P* = 0.002].

### Experiment 2. Effect of 2,5-AM on hepatic energy metabolism in OP and OR rats

At the time of 2,5-AM injection, mean body weights of rats identified as OP and OR rats were identical (257 ± 6 and 257 ± 3 g, respectively). One hour after 2,5-AM injection, hepatic ATP content (Figure 4) was lower in OP rats than it was in OR rats, but not significantly so [*t*(14) = 1.63, P = 0.12], while ATP/ADP ratio and phosphorylation potential were significantly lower in OP rats compared with OR rats [*t*(14) = 2.6, *P* < 0.021 and *t*(14) = 2.2, *P* < 0.043, respectively].

**Figure 4.**
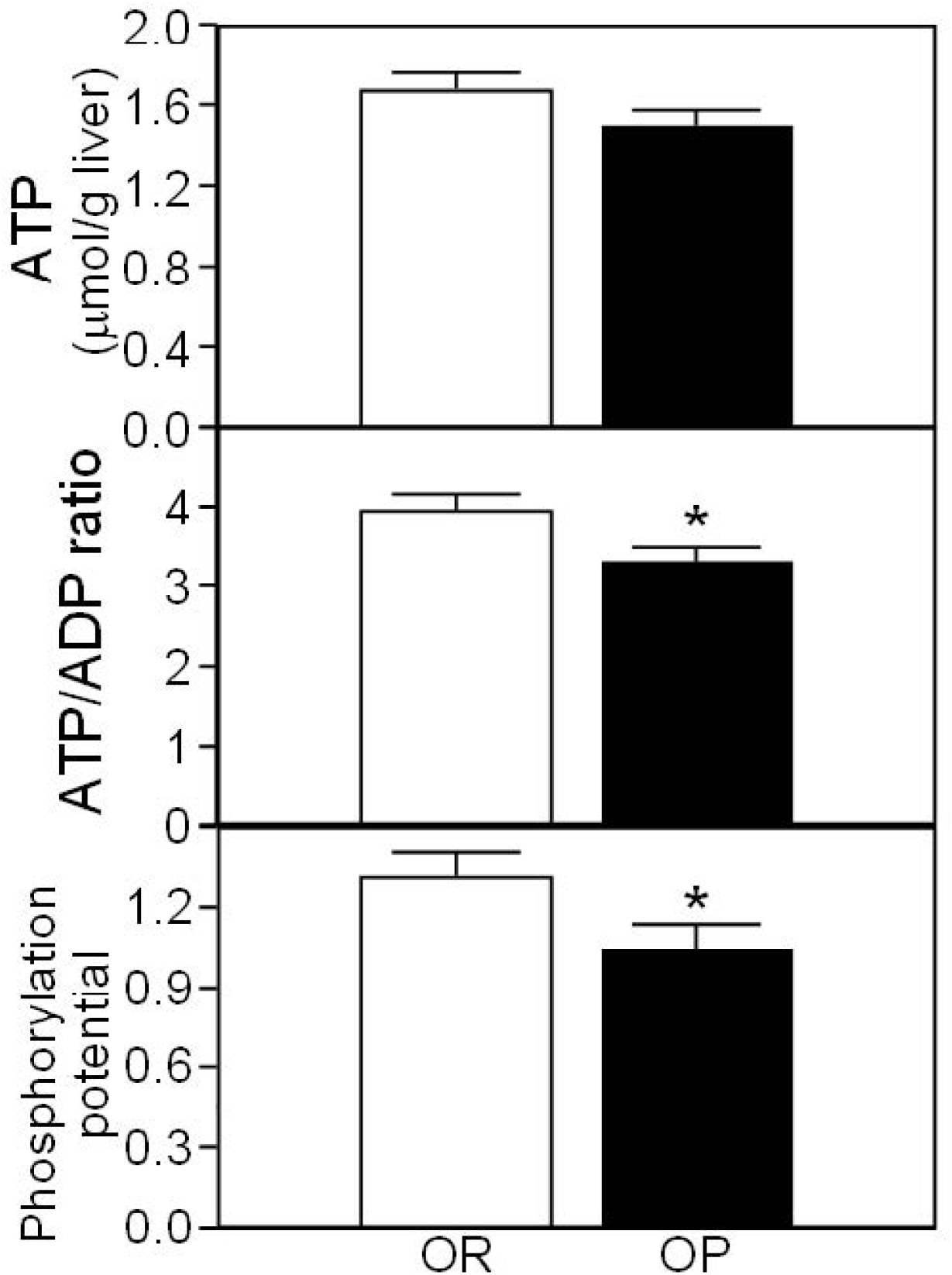
Liver energy status 1 h after injection of 2,5-AM (300 mg/kg body weight., i.p.) in obesity-prone (OP) and obesity-resistant (OR) rats. Values are mean ± SEM. N = 8 per group. * denotes a significant difference (P < 0.05; t-test) between groups.

## DISCUSSION

These results suggest that susceptibility to diet-induced obesity is associated with a limited capacity to defend liver energy status. Compared with OR rats, OP rats had lower hepatic energy status after a fast as measured by ATP:ADP ratio and phosphorylation potential. Restoration of liver energy status during subsequent refeeding was delayed in OP rats assessed by these measures as well as by hepatic ATP concentrations. We previously reported (21) that the time course of compensatory hyperphagia during refeeding after a fast paralleled that of the restoration of liver energy status in outbred rats. In the experiment reported here, outbred OP rats consumed more food during refeeding than did OR rats and differences between these two groups in the time course of eating behavior paralleled in an inverse fashion those in energy status when differences in food consumption were eliminated by restricting food intake. These findings are consistent with the hypothesis that decreased hepatic energy status is a stimulus for eating behavior (23,24) and suggest that increased food intake in diet-induced obesity is driven by such a signal.

OP and OR rats differed with respect to other metabolic parameters measured after fasting and during refeeding, but none appeared to account for the differences in compensatory intakes between the two groups. OP rats had greater plasma glucagon and leptin concentrations during refeeding than did OR rats, but changes in the plasma level of these hormones over time did not faithfully correspond to those in food intake during refeeding. Administration of glucagon and leptin are typically associated with satiety or reduction of food intake, further suggesting that increases in the circulating concentrations of these hormones were not responsible for the greater intakes of OP rats during refeeding.

Plasma FFA concentrations were greater in OP than in OR rats after fasting and by 3 h during refeeding at which time, as expected, they had decreased markedly in both groups. Plasma triglycerides concentrations increased three-fold from fasting levels in OP rats by 3 h of refeeding, compared to a 45% increase in OR rats, and then fell by 6 h during refeeding to levels comparable to those in OR rats and after fasting. Further studies would be needed to determine the cause of the spike in circulating triglycerides in OP rats, although more rapid clearance of fat from the gastrointestinal tract, reduced clearance of triglycerides from the circulation, and/or greater rates of lipogenesis or re-esterification of plasma fatty acids could be involved.

Although in Experiment 1 rats were switched to the HC/LF diet after screening for OP/OR status by feeding the HC/HF diet, the body weights of OP rats did not return to levels seen in OR rats by the time of the fasting-refeeding tests. This limits interpretation of the results because it is not clear whether the differences between OP and OR rats in food intake, liver energy status or other metabolic parameters were due to those in susceptibility to obesity or to differences in body fat content or its consequences. However, the experiment with 2,5-AM did not suffer from this potential confound due to differences in body weight, thus suggesting that at least in this case, the greater increase in food intake observed in OP rats during refeeding is more easily associated with liver energy status than fat oxidation.

We have reported previously (9) that outbred rats like those used in the present experiment can be screened for OP/OR status without feeding a HC/HF diet that creates potentially confounding effects of ongoing or previous differences in body weights of the two groups. We used this screening method in Experiment 2 to avoid the confounding effects of weight differences between OP and OR rats to explore in a preliminary fashion whether liver energy status in OP rats is more vulnerable to injection of 2,5-AM, which has been shown to reduce liver energy status in rats (11). All three measures of liver energy status were lower in OP rats after injection of 2,5-AM, with ATP:ADP ratio and phosphorylation index significantly so. Because liver energy status was not determined under a control (saline injection) condition, it is not clear whether this result reflects a greater vulnerability to a reduction of liver energy status or a lower basal energy status in OP rats. The observation that liver energy status in OP and OR rats was similar after 12 h of refeeding might suggest that it would also be comparable under the ad libitum feeding condition used for the 2,5-AM experiment. Additional studies are needed to clarify this issue.

Recognizing the limitations of the experiments described here, the results are consistent with studies reporting reduced liver energy status in diet-induced and genetically obese animals (25–28) and suggest that such a deficit in liver energy metabolism may underlie the susceptibility to diet-induced obesity. Co-administration of a fatty acid oxidation inhibitor and 2,5-AM, which in doses that are without effect alone, synergistically decreases liver energy status and increases food intake (19). A greater vulnerability of OP rats to the effects of 2,5-AM on liver energy status would be consistent with such a synergistic response inasmuch as these animals have a preexisting, limited capacity for hepatic fatty acid oxidation (6–9). Fructose also suppresses liver energy status (29,30), including in humans consuming a fructose solution (31). Excessive fructose consumption is considered a risk factor for human obesity (32,33). Considering that humans prone to obesity also have a reduced capacity to oxidize fat (e.g., 33,34), it is tempting to speculate that fructose consumption poses an even greater risk for weight gain in such people.

## Acknowledgements

This work was supported by NIH grant DK-53109.

## References

1. Ramirez I, Friedman MI. Dietary hyperphagia in rats: Role of fat, carbohydrate, and energy content. Physiol Behav 1990;47:1157–63.

2. Tordoff MG, Ellis HT. Obesity in C57BL/6J mice fed diets differing in carbohydrate and fat but not energy content. Physiol Behav 2022;243:113644.

3. Levin BE, Triscari J, Sullivan AC. Relationship between sympathetic activity and diet-induced obesity in two rat strains. Am J Physiol 1983;245:R364–71.

4. Levin BE, Dunn-Meynell AA, Balkan B, Keesey RE. Selective breeding for diet-induced obesity and resistance in Sprague-Dawley rats. Am J Physiol 1997;273:R725–30.

5. Burcelin R, Crivelli V, Dacosta A, Roy-Tirelli A, Thorens B. Heterogeneous metabolic adaptation of C57BL/6J mice to high-fat diet. Am J Physiol Endocrinol Metab 2002;282:E834–42.

6. Ji H, Friedman MI. Fasting plasma triglyceride levels and fat oxidation predict dietary obesity in rats. Physiol Behav 2003;78:767–72.

7. Ji H, Outterbridge LV, Friedman MI. Phenotype-based treatment of dietary obesity: differential effects of fenofibrate in obesity-prone and obesity-resistant rats. Metabolism 2005;54:421–9.

8. Ji H, Friedman MI. Reduced capacity for fatty acid oxidation in rats with inherited susceptibility to diet-induced obesity. Metabolism 2007;56:1124–30.

9. Ji H, Friedman MI. Reduced hepatocyte fatty acid oxidation in outbred rats prescreened for susceptibility to diet-induced obesity. Int J Obes 2008;32:1331–4.

10. Friedman MI, Harris RB, Ji H, Ramirez I, Tordoff MG. Fatty acid oxidation affects food intake by altering hepatic energy status. Am J Physiol 1999;276:R1046–53.

11. Friedman MI. An energy sensor for control of energy intake. Proc Nutr Soc 1997;56:41–50.

12. Rawson NE, Blum H, Osbakken MD, Friedman MI. Hepatic phosphate trapping, decreased ATP, and increased feeding after 2,5-anhydro-D-mannitol. Am J Physiol 1994;266:R112–7.

13. Tordoff MG, Rawson N, Friedman MI. 2,5-anhydro-D-mannitol acts in liver to initiate feeding. Am J Physiol 1991;261:R283–8.

14. Rawson NE, Friedman MI. Phosphate loading prevents the decrease in ATP and increase in food intake produced by 2,5-anhydro-D-mannitol. Am J Physiol 1994;266:R1792–6.

15. Koch JE, Ji H, Osbakken MD, Friedman MI. Temporal relationships between eating behavior and liver adenine nucleotides in rats treated with 2,5-AM. Am J Physiol 1998;274:R610–7.

16. Park CR, Seeley RJ, Benthem L, Friedman MI, Woods SC. Whole body energy expenditure and fuel oxidation after 2,5-anhydro-D-mannitol administration. Am J Physiol 1995;268:R299–302.

17. Ji H, Graczyk-Milbrandt G, Osbakken MD, Friedman MI. Interactions of dietary fat and 2,5-anhydro-D-mannitol on energy metabolism in isolated rat hepatocytes. Am J Physiol Regul Integr Comp Physiol 2002;282:R715–20.

18. Friedman MI, Koch JE, Graczyk-Milbrandt G, Ulrich PM, Osbakken MD. High-fat diet prevents eating response and attenuates liver ATP decline in rats given 2,5-anhydro-D -mannitol. Am J Physiol Regul Integr Comp Physiol 2002;282:R710–4.

19. Ji H, Graczyk-Milbrandt G, Friedman MI. Metabolic inhibitors synergistically decrease hepatic energy status and increase food intake. Am J Physiol Regul Integr Comp Physiol 2000;278:R1579–82.

20. Start C, Newsholme EA. The effects of starvation and alloxan-diabetes on the contents of citrate and other metabolic intermediates in rat liver. Biochem J 1968;107:411–5.

21. Ji H, Friedman M. Compensatory hyperphagia after fasting tracks recovery of liver energy status. Physiology & Behavior 1999;68:181–6.

22. Ramirez I. Physiological and biochemical measurements in relation to feeding. In: Toates FM, Rowland NE, editors. Feeding and Drinking. Amsterdam: Elsevier; 1987. p. 151–65.

23. Friedman M. Food intake: control, regulation, and the illusion of dysregulation. In: Harris R, Mattes R, editors. Appetite and food intake: behavioral and physiological considerations. Boca Raton, FL: CRC Press; 2008. p. 1–19.

24. Watts AG, Kanoski SE, Sanchez-Watts G, Langhans W. The physiological control of eating: signals, neurons, and networks. Physiological Reviews 2022;102:689–813.

25. Chavin KD, Yang S, Lin HZ, Chatham J, Chacko VP, Hoek JB, Walajtys-Rode E, Rashid A, Chen C-H, Huang C-C, et al. Obesity induces expression of uncoupling protein-2 in hepatocytes and promotes liver ATP depletion. J Biol Chem 1999;274:5692–700.

26. Serkova NJ, Jackman M, Brown JL, Liu T, Hirose R, Roberts JP, Maher JJ, Niemann CU. Metabolic profiling of livers and blood from obese Zucker rats. J Hepatol 2006;44:956–62.

27. López-Soldado I, Zafra D, Duran J, Adrover A, Calbó J, Guinovart JJ. Liver glycogen reduces food intake and attenuates obesity in a high-fat diet–fed mouse model. Diabetes 2015;64:796–807.

28. Geisler CE, Ghimire S, Hepler C, Miller KE, Bruggink SM, Kentch KP, Higgins MR, Banek CT, Yoshino J, Klein S, et al. Hepatocyte membrane potential regulates serum insulin and insulin sensitivity by altering hepatic GABA release. Cell Rep 2021;35:109298.

29. Oberhaensli RD, Galloway GJ, Taylor DJ, Bore PJ, Radda GK. Assessment of human liver metabolism by phosphorus-31 magnetic resonance spectroscopy. BJR 1986;59:695–9.

30. Raivio KO, Kekomäki MP, Mäenpää PH. Depletion of liver adenine nucleotides induced by D-fructose. Dose-dependence and specificity of the fructose effect. Biochem Pharmacol 1969;18:2615–24.

31. Bawden SJ, Stephenson MC, Ciampi E, Hunter K, Marciani L, Macdonald IA, Aithal GP, Morris PG, Gowland PA. Investigating the effects of an oral fructose challenge on hepatic ATP reserves in healthy volunteers: A 31P MRS study. Clin Nutr 2016;35:645–9.

32. Tappy L. Fructose-containing caloric sweeteners as a cause of obesity and metabolic disorders. J Exp Biol 2018;221:jeb164202.

33. Herman MA, Birnbaum MJ. Molecular aspects of fructose metabolism and metabolic disease. Cell Metab 2021;33:2329–54.

34. Raben A, Andersen HB, Christensen NJ, Madsen J, Holst JJ, Astrup A. Evidence for an abnormal postprandial response to a high-fat meal in women predisposed to obesity. American Journal of Physiology-Endocrinology and Metabolism 1994;267:E549–59.

35. Filozof CM, Murúa C, Sanchez MP, Brailovsky C, Perman M, Gonzalez CD, Ravussin E. Low plasma leptin concentration and low rates of fat oxidation in weight-stable post-obese subjects. Obesity Research 2000;8:205–10.

